# A spatial approach to jointly estimate Wright’s neighborhood size and long-term effective population size

**DOI:** 10.1101/2023.03.10.532094

**Authors:** Zachary B. Hancock, Rachel H. Toczydlowski, Gideon S. Bradburd

## Abstract

Spatially continuous patterns of genetic differentiation, which are common in nature, are often poorly described by existing population genetic theory or methods that assume panmixia or discrete, clearly definable populations. There is therefore a need for statistical approaches in population genetics that can accommodate continuous geographic structure, and that ideally use georeferenced individuals as the unit of analysis, rather than populations or subpopulations. In addition, researchers are often interested describing the diversity of a population distributed continuously in space, and this diversity is intimately linked to the dispersal potential of the organism. A statistical model that leverages information from patterns of isolation-by-distance to jointly infer parameters that control local demography (such as Wright’s neighborhood size), and the long-term effective size (*N_e_*) of a population would be useful. Here, we introduce such a model that uses individual-level pairwise genetic and geographic distances to infer Wright’s neighborhood size and long-term *N_e_*. We demonstrate the utility of our model by applying it to complex, forward-time demographic simulations as well as an empirical dataset of the Red Sea clownfish (*Amphiprion bicinctus*). The model performed well on simulated data relative to alternative approaches and produced reasonable empirical results given the natural history of clownfish. The resulting inferences provide important insights into the population genetic dynamics of spatially structure populations.

## Introduction

In many species, individual (or gamete) dispersal is geographically limited, leading to spatial structure. This spatial structure can in turn give rise to a pattern of isolation by distance (Wright, 1943, 1946; Meirmans 2012), in which a focal individual is, on average, more closely related to an individual sampled nearby than it is to another individual sampled farther away. Much of early population genetic theory was derived under the assumption of random mating, which may have adequately described the model organism populations common in empirical population genetic studies of the time (e.g., *Drosophila* in vials; Prout 1954; Merrell 1953; Dobzhanksy & Spassky 1962) but is less well-suited to describing spatially continuous genetic structure. While several theoretical approaches have relaxed these assumptions by modeling the population as partitioned into “demes” with some constant rate of migration between them (e.g., Wright’s [1943] “island model” and Kimura & Weiss’ [1964] “stepping-stone model”), each approach still maintained panmixia within demes, and neither effectively captures population genetic dynamics in continuous space. The island and stepping-stone models inspired a series of statistical approaches that rely on partitioning samples into discrete populations with some level of genetic differentiation (sometimes estimated) between them (e.g., Wright 1951; Pritchard et al. 2000; Pickrell & Pritchard 2012; Peter 2016). These approaches have been expanded to estimate the degree of admixture between these discrete populations (e.g., ADMIXTURE – Alexander et al. 2009), where individuals inferred to have ancestry proportions in multiple inferred clusters are described as “admixed.” However, when the true pattern of genetic variation is continuous across the landscape, the inferred *K* clusters in a method like STRUCTURE or individual admixture proportions in ADMIXTURE are mere artifacts of the sampling scheme (Frantz et al 2009, Bradburd et al 2018). An example of this phenomenon in action can be found in studies that group human genetic variation by continent, which often generates an apparent pattern of discrete clusters (Rosenberg et al. 2002; Li et al. 2008); however, when *individuals* are the unit of investigation, human genetic ancestry is continuous and defies simple continental or population groupings (Ramachandran et al. 2005; Lewis et al. 2022; Carlson et al. 2022).

Wright (1943) and Malécot (1946) were the first to consider population models in which individuals were continuously distributed across one- and two-dimensions in geographic space. Wright (1943, 1946) introduced the concept of “neighborhood size” 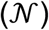 as a statistic to describe natural populations; 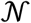 was meant to capture the number of potential parents within a given radius of a focal individual, where that radius was defined by the dispersal distance in two-dimensions. Their early theoretical work has since been expanded by Malécot (1948), Maruyama (1971), Nagylaki (1975; 1978), Felsenstein (1976), Barton et al. (2002), Barton et al. (2013), among others. Despite these theoretical advances, relatively few statistical methods exist for examining populations in continuous space. Wright’s neighborhood size 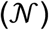 provides important information about the dispersal potential and the rate of genetic drift in continuously distributed populations at a localized level, and is therefore a useful quantity to know, both for conservation purposes and a general understanding of the evolutionary context of a particular population or species.

Rousset (1997; 2000) introduced a method for the estimation of Wright’s neighborhood size as the inverse of the slope of a regression between pairwise *F_ST_* / (1 – *F_ST_*) and the logarithm of pairwise geographic distance. While this method is limited in that it assumes a constant population density, it has the useful benefit of providing a single estimate of 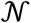 for the entire population (Shirk & Cushman 2014). Many popular programs enable researchers to estimate 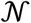 by implementing this expected relationship (e.g., SPAGeDI – Hardy & Vekemans 2002; Rousset & Leblois 2011). Importantly, the decision of which measure of *F_ST_* to use is non-trivial and can produce dramatically different results (Pearse & Crandall 2004; Bhatia et al. 2013). Furthermore, researchers must decide to estimate *F_ST_* between individuals or artificially designated subpopulations. The former is very sensitive to individual measures of genetic diversity, which can be particularly noisy and impacted by bioinformatic decisions that affect the presence of missing data and rare variants (Bhatia et al. 2013); this noisiness can be smoothed by lumping individuals into subpopulations, but this “lumping” approach is not ideal when sampling covers a large geographic area and there is a continuous pattern of isolation-by-distance (Pearse & Crandall 2004). An alternative method, introduced by Shirk & Cushman (2014) and implemented in sGD (Shirk & Cushman 2011), estimates 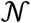 following a user-specified neighborhood radius, and then used the Burrow’s method of linkage disequilibrium (Cockerham & Weir 1977) to estimate the number of breeding individuals of the sample. However, this method does not utilize the information from the shape of the isolation by distance curve like Rousset’s. Furthermore, it relies on the investigator having strong prior knowledge on the dispersal potential of the organism such that they can adequately predict how to discretize space and, thus, how many samples should be included within the neighborhood estimation.

Another quantity that, like Wright’s neighborhood size, is useful for understanding and conserving species is the effective population size (*N_e_*). In a conservation setting, *N_e_* may serve as a rough proxy for census size and the adaptive potential of threatened populations (Exposito-Alonso et al. 2022; Theodoridis et al. 2021). Others have used *N_e_* to investigate the relationship between range size or dispersal ability (Leigh et al. 2021; De Kort et al. 2021). Estimation of the inbreeding effective size *N_e_* (Wright 1931) often relies on its relationship with genetic diversity, which, when the mutation rate is μ and the population is at mutation-drift equilibrium, is given by 4*N_e_μ* (termed the “population mutation rate”; Kimura & Crow 1964). This definition is distinct from the variance *N_e_*, which describes the rate of genetic drift between successive generations (Crow & Kimura 1970; Wang & Caballero 1999; Charlesworth 2009). A common estimator of the population mutation rate is Watterson’s *θ_w_*, which is *K / a_n_* where *K* is the number of segregating sites in the sample and 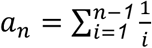 (Watterson 1975). However, Watterson’s *θ_w_* is naive to the spatial structure of the sample and is known to be upwardly biased relative to random mating expectations when neighborhood sizes are very low (Battey et al. 2020). Another estimator of the population mutation rate is 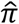, which is 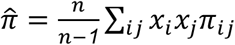, where *n* is the number of samples, *x_i_* and *x_j_* are the frequencies of the *i*th and *j*th sequence, and π*_ij_* is the number of nucleotide differences between the *i*th and *j*th site (Nei & Tajima 1981). At mutation-drift equilibrium and assuming no selection, 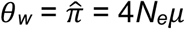.

In continuously distributed populations, inbreeding *N_e_* is not independent of 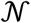 as each is impacted by dispersal (Barton et al. 2002; Wilkins 2004). When dispersal is high, *N_e_* converges to the population census size (*N_c_*) but tends to be much larger when dispersal is low. Similarly, because dispersal dictates the radius of 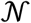, higher rates lead to more potential parents being included within 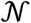. An ideal model of spatial population structure would thus be individual-based such that researchers would not need to arbitrarily group individuals into subpopulations and would co-estimate Wright’s neighborhood size and inbreeding *N_e_* while explicitly accounting for the shared influence of dispersal across timescales.

In this paper, we take steps towards this goal by introducing a model that jointly estimates 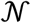 and long-term *N_e_* from data on pairwise π and geographic distance between individuals. We validate the model’s performance using individual-based forward-time simulations and evaluate our model’s performance against Rousset’s (1997) method when *F_ST_* is estimated between individuals. Finally, we apply our model to an empirical dataset of Red Sea clownfish (*Amphiprion bicinctus*; Saenz-Agudelo et al. 2015).

## Methods

### Model Intuition

The population genetic pedigree is ultimately shaped by an organism’s life-history, including its dispersal potential, generational structure, and mating strategies. Because mutations occur along the branches of the genetic genealogy contained within the population pedigree, the shape of the pedigree fundamentally determines the diversity of a sample as well as a whole host of additional summary statistics (Fig. 1). The field of statistical population genetics is predicated on the idea that information about processes shaping the population pedigree (e.g., selection, demography) leave their imprint in patterns of genetic diversity and divergence observable in a modern-day sample.

**Figure 1.**
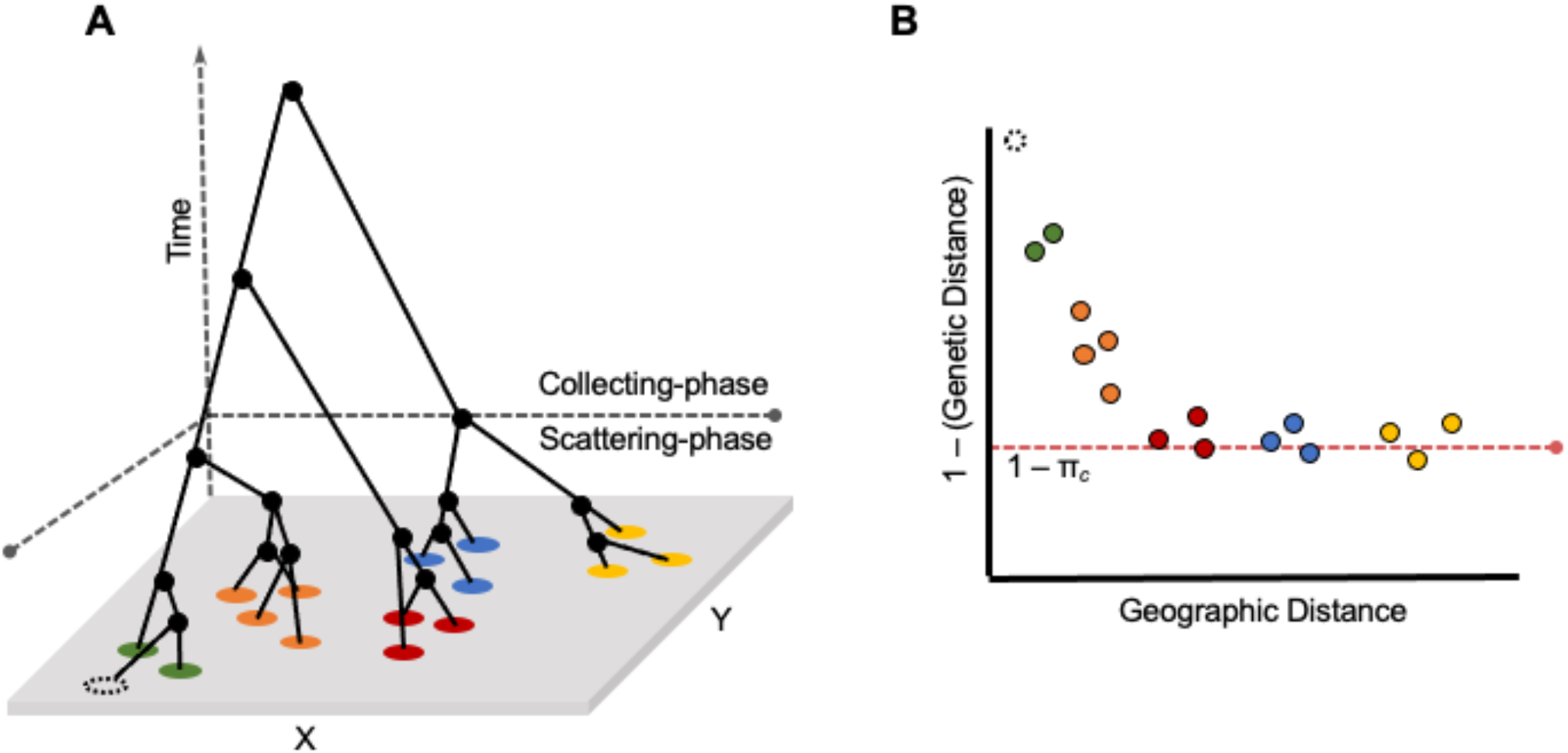
Relationship between the separation-of-timescales of the coalescent and the isolation-by-distance (IBD) curve in continuous space. A) A population genetic tree showing the relationships between sampled individuals (colored circles) across a continuous landscape with dimensionality (*x*, *y*). Relative to a focal sample (dotted circle), the transition to the collecting phase occurs as the rate of coalescence converges to a neutral Kingman’s coalescent. B) An IBD plot relative to the focal individual. The transition to the collecting phase occurs when geographic distance is no longer predictive of genetic distance. The red dotted line denotes *s* (or 1 – π_*c*_), which is the estimated mean minimum relatedness between individuals in the population.

For populations that are spatially structured such that there is autocorrelation between geographic location and genetic ancestry, the pedigree becomes distorted relative to random-mating expectations. This pedigree distortion occurs in two phases: the *scattering phase* and the *collecting phase* (Wakeley 1999, Wilkins 2004; Wilkins & Wakeley 2002). Going backward in time from the present, the scattering phase happens first, and is characterized by lineages that are geographically near one another coalescing more rapidly than expected under random mating (i.e., with a probability greater than 1 / 2*N_e_*; Wilkins 2004; Wilkins & Wakeley 2002). This signature is especially strong when dispersal is low, as nearby individuals are more likely to be more closely related than a pair of individuals selected from the population at random (with respect to geography). The parameter that governs the rate of coalescence in this phase of the population pedigree is Wright’s neighborhood size, which is defined as 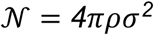, where π is the mathematical constant, *ρ* is population density, and *σ* is the standard deviation of the effective dispersal distance defined by a normal distribution with mean zero (Wright 1940, 1943, 1946, 1949). Forwards-in-time, 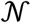 describes the number of potential mates within a circle of radius 2*σ*^2^, within which breeding occurs approximately at random with respect to geographic position. Backwards-in-time, Wright’s neighborhood size can be thought of as the “pool of possible parents” of a focal individual (i.e., the number of reproductively mature individuals within two effective dispersal distances of the focal individual in any direction).

Further in the past — exactly how far depends on the rate of dispersal and the habitat geometry (Wilkins 2004) — the coalescent process shifts to the second phase, known as the *collecting phase*, in which the rate of coalescence is independent of the geographic distribution of the modern-day sample of individuals. This independence arises because, following a focal individual’s pedigree backward through time and across space, the geographic distribution of its genetic ancestors expands until it ceases to be correlated with that focal individual’s location (Fig. 1b; Bradburd & Ralph 2019). In the collecting phase, relatedness between lineages is well-represented by a neutral coalescent process (Kingman 1982), in which the rate of coalescence per generation is 1 / 2*N_e_* and the average time to the most recent common ancestor is 4*N_e_* generations.

This two-phase distortion of the shape of the pedigree relative to random-mating expectations affects estimates of the genetic diversity of the population. In a spatially structured population at migration-drift equilibrium, dispersal limitations shrink the depth of the coalescent tree locally (i.e., individuals nearby are more related on average than expected under panmixia) while expanding it globally (farther away, individuals are more distantly related than expected). As a result, estimates of 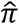 or 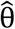 are highly dependent on the spatial scale of sampling, and are generally downwardly biased relative to the equilibrium π in the collecting-phase; this effect is particularly strong when dispersal is very low relative to the length of the range and the geographic area encompassed by the genotyped samples is small (Exposito-Alonso et al 2022).

Our model (explained below) relies on the relationship between the spatial population genetic pedigree and the isolation-by-distance curve (Fig. 1b, 2). At short geographic distances, there is strong spatial autocorrelation of relatedness - this captures the scattering-phase of the spatial pedigree, and it decays rapidly (Fig. 1b). As the curve flattens, geographic distance ceases to be explanatory of relatedness and the population approaches expectations under panmixia - this is capturing the transition to the collecting-phase. The shape of the decay of relatedness over short spatial scales carries information about 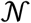, whereas the inferred asymptote of relatedness over large geographic distances represents a diversity equilibrium (what we term “collecting-phase π”, π_c_), and is most informative about long-term inbreeding *N_e_*.

**Figure 2.**
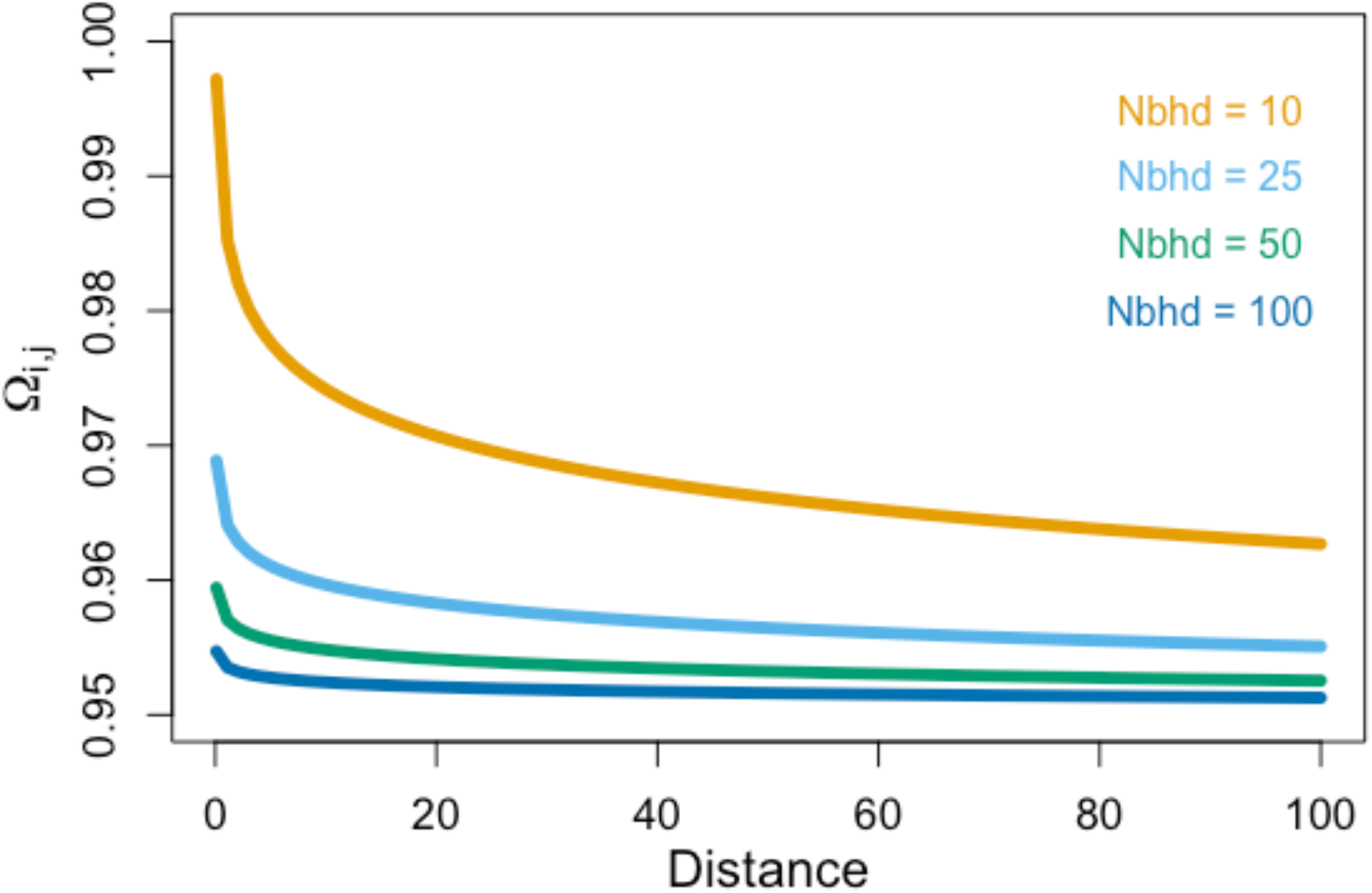
Expected relatedness decay curves of *Ω_i,j_* with distance given *s* = 0.95 for various values of neighborhood size (“Nbhd”). See Equation 5 in the text.

### Model

We first introduce the model of isolation-by-distance (IBD) that we use - both its form and its assumptions, then describe how we fit this model to observed genomic data. Briefly, our model is closely related to previous theoretical models of genetic differentiation in continuous space (e.g., Wright 1943; Malécot 1946; Barton et al. 2002, 2010, 2013; Ringbauer et al. 2017), and describes the decay in pairwise homozygosity with the geographic distance between samples assuming a homogeneous landscape with isotropic dispersal. We implement this model in a Bayesian framework to estimate the posterior distribution of model parameters conditioned on observed pairwise sample homozygosity and pairwise geographic distance between samples.

Our model seeks to capture two important components of spatially structured populations: 1) that samples covary in their allele frequencies, with a covariance that decays with geographic distance during the scattering phase, and 2) that there exists an equilibrium level of minimum divergence between individuals that is established during the collecting phase (Fig. 1). Furthermore, our model assumes that populations exist in continuous space in two-dimensions and that dispersal is random and diffusive. In a single dimension, tracking diffusive dispersal backwards-in-time, two lineages will eventually exist in the same location at the same time at some point in the past. However, in two-dimensions, lineages diffusing via Brownian motion will never arrive in the same place at the same time (e.g., Nagylaki 1978, Barton et al. 2002). Modeling a spatial coalescent process in two dimensions is therefore tricky. This issue has been circumvented in the past (Wright 1943; Malécot 1946) by assuming that individuals need not be in the exact same location at the same time, but merely within a given radius of one another. Within this radius, individuals are assumed to interact in a way that is independent of the geographic distance between them, so that the probability of coalescence can be described as 1 / 2*ρ* (Barton et al. 2002), where *ρ* is population density. For habitats that are relatively homogenous, such that geographic distance is the primary explanation of covariance, with constant *ρ* and σ through time and across space, the probability of samples *i* and *j* being identical-by-descent (F_ij_) can be estimated by

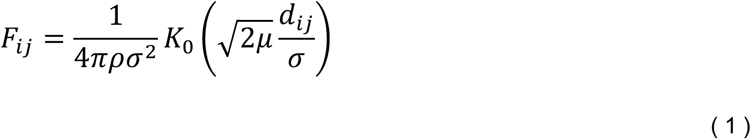

where *K*_0_ is the modified Bessel function of the second kind of order 0, *d_ij_* is the pairwise geographic distance between samples *i* and *j*, and π is the mathematical constant (Wright 1943, Malécot 1946, Barton et al. 2002). Barton et al. (2002) notes that this approximation diverges as *d_ij_* → *0*; to account for this, following Ringbauer et al. (2017) we designate a short distance, *κ*, within which the rate of coalescence becomes a constant *γ* that no longer depends on the geographic distance between samples. We refer to *γ* as the “in-deme” rate of coalescence. Theoretically, *γ* should converge on 1 / 2*ρ* and an optimal value for *κ* would be approximately 2*σ* (Barton et al. 2002).

The classic Wright-Malécot formula (Eqn. 1) describes the probability of identity-by-descent, which, in an infinite population, theoretically decays to zero. However, our model breaks the assumptions of the Wright-Malécot model in two important ways. First, we assume that populations are finite, meaning that there will be a maximum time at which all individuals in the population are identical-by-descent. Second, we choose to model identity-by-state, rather than identity-by-descent, as we assume more empiricists will have access to identity-by-state information than identity-by-descent (particularly in non-model organisms). Therefore, we must incorporate into our model a background rate of genetic similarity at which all individuals in the population are identical-by-state. The expected homozygosity of a pair of samples, *i* and *j*, is thus

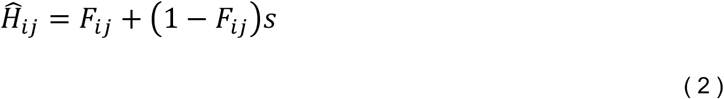

where 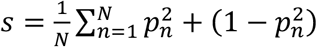 and *p_n_* is the population frequency of an allele at the *n*th of *N* loci (Ringbauer et al. 2018). The quantity *s* represents the “background” rate of sequence similarity and can be thought of as the complement of the amount of genetic diversity in a population at equilibrium: 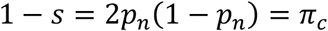, which we define as “collecting-phase π” and is related to long-term *N_e_*. Unlike 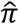 (mean pairwise genetic distance in a population, Nei & Tajima 1981), estimates of *π_c_* (defined by the asymptote of the IBD curve, which showcases the transition to the collecting phase; Fig. 1) should be insensitive to the size of a sampled area.

### Inference and Implementation

We assume users’ data consist of allele frequencies taken across *L* unlinked, biallelic single nucleotide polymorphisms (SNPs) genotyped across a set of *N* samples. Each sample may consist of a single individual or a group of individuals collected at a single location. Allele frequencies may be estimated from genotype data (e.g., the frequency of the *l*th allele in the *n*th sample is simply the number of times that allele is observed divided by the total number of genotyped haplotypes in that sample at that locus) or from pooled sequencing data. From these data, we compute the sample pairwise homozygosity, which is the complement of the pairwise diversity between the samples (i.e., 1 – *D_xy_*). We note that users working with low-coverage sequence data may wish to generate estimates of *D_xy_* without conditioning on allele frequencies (e.g., Buerkle & Gompert 2012; Ellegren 2014). Pairwise homozygosity between samples *i* and *j* gives the probability that, at a locus chosen at random, a pair of alleles sampled at random from *i* and *j* respectively, are the same. We calculate it as:

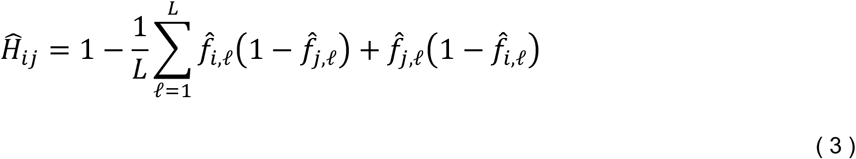

where, 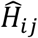 gives the sample homozygosity between samples *i* and *j* calculated across all *L* loci, and 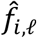 gives the sample allele frequency in the *i*th sample at the *l*th locus. Pairwise homozygosity is a measure of absolute genetic similarity, so it is not sensitive to the sampling configuration. Additionally, 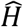 is proportional to the allelic diversity defined in Bradburd et al. (2018), so we proceed by assuming it can be reasonably modeled as Wishart-distributed, and the framework we use for statistical inference is similar to that of Bradburd et al (2018) (see also Ringbauer et al [2018]).

Specifically, we construct a parametric expected homozygosity matrix using a modified version of the Wright-Malécot model of isolation by distance introduced in Equation 2 and calculate the likelihood of the sample homozygosity as a draw from a Wishart distribution parameterized by the parametric homozygosity. Concretely, we write that the probability of identity by descent between samples *i* and *j, F_ij_*, is

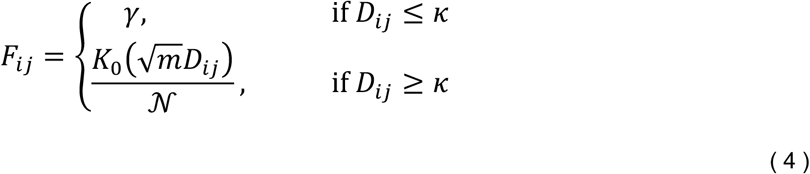

where γ is the “in-deme” rate of identity by descent (i.e., the probability of being identical by descent at distances short enough that mating can reasonably be considered panmictic, *D_ij_* ≤ *κ*), and *F_ij_* for *D_ij_* ≥ *κ* is given by the Wright-Malécot function introduced in Equation 2. We then construct our parametric homozygosity between samples *i* and *j*, Ω*_ij_*, as

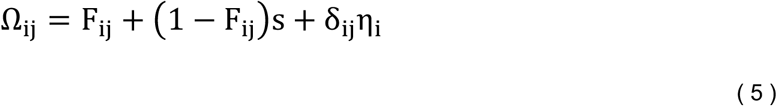

where *s* is the rate of identity by state not due to identity by descent, δ_ij_ is the Kronecker δ, and η_i_ is a parameter (often called a “nugget” in the geostatistical literature, Diggle et al 1998) that captures inbreeding specific to the *i*th sample. We then calculate our likelihood as

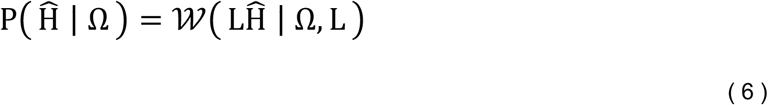

where *L* is the number of independent genomic loci used in the calculation of 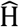.

We take a Bayesian approach to infer the parameters of this model. The posterior probability density of our parameters is given by

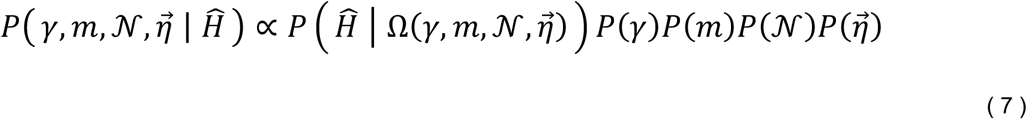

where 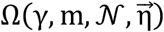 denotes the dependence of Ω on its constituent parameters 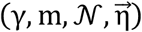, and *P*(*θ*) denotes the prior probability of a given parameter θ. Table S1 describes the prior probability distributions we implement for each parameter in our model. We implement this model in Rstan (Stan Development Team 2023) and use STAN’s Hamiltonian Monte Carlo algorithm to characterize the posterior distribution of the parameters.

Because pairwise homozygosity often varies over a very small absolute range (e.g., 0.99-0.999), we take several steps to facilitate inference on the parameters of the model. First, we estimate the parameters *m*, γ, and 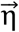 in log space, which should help chains mix over the posterior density. Second, we scale both the sample homozygosity 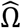 and the parametric homozygosity Ω to vary between 0 and 1:

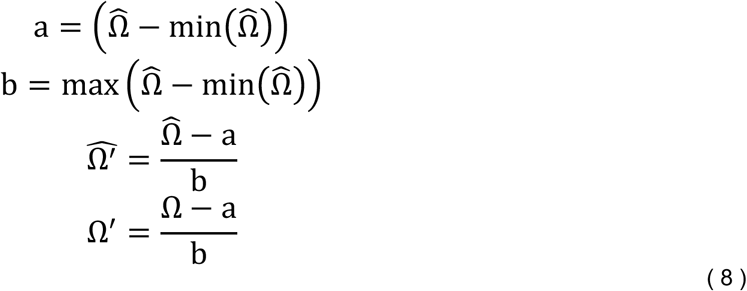

This scaling is not without drawbacks, as the variance of a Wishart distribution parameterized by Ω is not the same as that parameterized by Ω’. However, we feel that the benefits it offers outweigh the costs, particularly because, to our knowledge, there is no theoretically motivated “correct” variance on homozygosity.

### Simulations

We evaluated model performance using individual-based forward-time simulations performed in SLiM v3.6 (Haller & Messer 2019). Simulated individuals were diploid (2*n*) and hermaphroditic, with haploid genomes of 100 Mb, non-overlapping generations, and a uniform recombination rate of 10^−9^ per base-pair per generation. Mate choice, spatial competition, and dispersal are controlled by a constant value, σ. Individuals were simulated on a continuous, two-dimensional 25×25 landscape with reflecting boundaries. Total population density was regulated by an enforced carrying-capacity, *K*, to avoid spatial clumping (Felsenstein 1975). To reduce the impact of edge effects, we designate this density-dependent competition using the function *localPopulationDensity(*) in SLiM, which computes the total interaction strength around a focal individual first as 2π*σ*^2^, then divides this strength by the integral of the interaction after clipping by the bounds of the specified landscape (Haller & Messer 2022). This has the desired effect of rescaling competition relative to the occupiable area. Edge-effects reduced the census population size by ~35% relative to *Kw*^2^, where *w* is the width of the simulated landscape. The maximum distance at which individuals experience spatial competition was capped at 3*σ*. Similar to spatial competition, mates were chosen within a maximum distance of 3σ, with each potential mate assigned a weight drawn from a Gaussian distribution with max 1 / 2π*σ*^2^. The number of offspring produced per mating event was drawn from a Poisson distribution with shape parameter *λ* = 2, which on average replaces the parents. Finally, offspring dispersal distance was drawn from a normal distribution with mean 0 and standard deviation *σ* and maximum of 3*σ*.

We performed simulations across a range of values of *K* (2, 5, 10, 25) and *σ* (0.5, 0.75, 1.0, 1.25, 1.5, 2.0), which varied theoretical neighborhood size from a minimum of 6.25 to a maximum of 861.81, and total census size from ~800–10,000. We performed 10 replicates for each combination of parameter values, for a total of 240 simulations. Each simulation was run for 100,000 generations to ensure time for dispersal to shape patterns of genetic diversity. The output from SLiM were tree-sequences and these were parsed in Python using the package *pyslim* v.1.0.1 (Kelleher et al. 2018). For trees in which multiple roots existed (i.e., coalescence had not yet occurred during the SLiM run), we performed “recapitation,” which simulates a neutral coalescent process among remaining, uncoalesced lineages (Haller et al. 2018). Mutations were then simulated onto the tree-sequences using *msprime* v.1.2.0 (Kelleher et al. 2016) at a rate of 10^−7^ per base-pair per generation. We then randomly sampled 100 individuals alive in the final generation and we calculated pairwise π between each pair of individuals using *tskit* v.0.5.3 (Kelleher et al. 2018). We output the pairwise individual π matrix and geographic coordinate matrix, the latter of which was converted into a distance matrix in R (R Core Team 2022) using the package *fields* (Nychka et al. 2021). These two matrices were then used as input to our model, which we used to infer a range of parameters (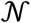, π_c_, *γ*, and *μ*). Note that of these parameters, we only explicitly designated *μ*; the remaining values have theoretical expectations given the parameters we chose for each individual simulation but are not explicitly defined. Theoretical *N_e_* in a square habitat is

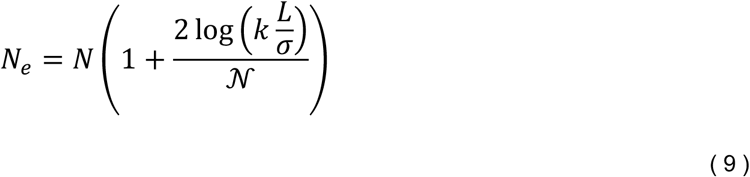

where *k* = 0.24 when dispersal is Gaussian, and *L* is the length of the major axis (Wilkins 2004; Barton et al. 2002; Charlesworth et al. 2003). Importantly, as the rate of dispersal approaches the length of the range and 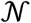 approaches the census size, *N_e_* approximately equals the population census size, *N*. However, theoretical *N_e_* is likely an overestimate of the true *N_e_* in our simulations due to reduced population density near the edges as opposed to the expected homogenous distribution. Thus, we estimate the “true” *N_e_* in each simulation as 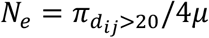, where 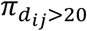 is the mean pairwise diversity for samples at distances greater than 20, ensuring they all occur in the collecting phase. For 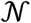, we do not use a similar proxy and instead rely on the theoretical 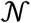 given that the impacts of deviations from theory in our simulations for areas of relevance to 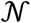 are expected to be much smaller than for *N_e_*.

For each dataset, we performed four MCMC chains of 4000 steps each. We pruned the first 1e3 steps in each chain as burn-in and thinned the remaining steps by sampling every 4^th^ iteration, for a total of 250 sampled post burn-in iterations per chain. Because some chains displayed poor mixing, we selected the chain with the highest mean posterior probability as the one from which to report results. We visually verified that the chains had achieved convergence by inspecting the trace plots of the posterior probability and parameter values of interest.

Simulations and model estimation were performed on the Michigan State Institute for Cyber-Enabled Research High Performance Computing Cluster (ICER HPCC). Code, including the ICER HPCC slurm workflow, SLiM recipes, and Python scripts, can be found at https://github.com/zachbhancock/WM_model.

### Empirical dataset

We used a reduced-representation genomic dataset of Red Sea clownfish (*Amphiprion bicinctus*) from Saenz-Agudelo et al. (2015) to explore how this model performed on empirical data. This dataset is composed of 103 wild individuals sampled from 10 unique locations spread across the species’ range (Red Sea). Sequence reads were generated using double-digest restriction-associated DNA and sequenced 1 × 101 base pairs. We downloaded the raw (demultiplexed) sequence reads for each of the 103 individuals from the International Nucleotide Sequence Database Collaboration (BioProject PRJNA294760, https://www.ncbi.nlm.nih.gov/bioproject/PRJNA294760) using the fasterq-dump function in the SRA Toolkit (V2.10.7, Kodama, Shumway, & Leinonen 2012). We dropped reads with uncalled bases and where the mean Phred score was < 15 (sliding window 15% of read length; Stacks2 process_radtags module V2.54 – Rochette, Rivera-Colón, & Catchen 2019). We confirmed that no common adapter sequences were present (3’ end of reads) and that all reads were a uniform length (81 bp post demultiplexing and trimming by Saenz-Agudelo et al.). Next, we assembled these reads *de novo* using Stacks2 (V2.54, Rochette, Rivera-Colón, & Catchen 2019) and optimized assembly parameters (see Saenz-Agudelo et al. 2015). Post Stacks, we dropped loci that were scored in <50% of individuals. Finally, we calculated pairwise pi and pairwise geographic distance between each pair of individuals in the dataset. Geographic distance here is measured as the distance traveling via the sea and avoiding crossing land (marmap – Pante & Simon-Bouhet 2013).

## Results

Our model performed well on both the simulated and empirical datasets. In the former, the model converged on the theoretical expectations for both 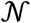 and *N_e_*. Furthermore, it performed favorably relative to Rousset’s method, which was unbiased but had much higher variance than our model, especially at lower 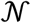. Finally, as applied to the empirical dataset, our model produced reasonable results given the known natural history of clownfish.

### Simulation results

The individual-based simulations were performed by varying both dispersal distance (*σ*) and local carrying-capacity (*K*), which generated a range of 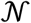 from 6.28 to 861.81. Given the simulated geographic range area, this range of values of encompasses 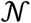 extreme/strong spatial structure that is likely never well-described by the Kingman’s coalescent at the lower end (Wilkins 2004) and virtually panmictic populations at the upper end.

At all simulated values of *σ* and *K*, the 95% equal-tailed credible interval of 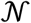 in the consistently included the theoretical 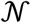; however, at larger *σ* and *K* it did so with high variance, with the median falling below the theoretical value (Fig. 3a). This is likely due to the reliance of the model on detectible signatures of covariance between geography and ancestry, which decays at large 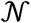 (Fig. 2). At small dispersal distances, the variance in our estimates of 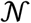 was relatively low even at higher *K*. However, at larger dispersal distance the variance subsequently increased regardless of *K*, indicating that *σ* likely has the largest impact on model precision. Notably, our model estimated an 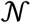 greater than the theoretical expectation only at the lowest *K* and *σ* combination, which may indicate that the Wright-Malécot model is a poor approximation of relatedness when 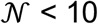 for a population of this geographic range with finite boundaries. Furthermore, we found that while Rousset’s method is an unbiased estimator, its variance becomes larger at lower *K* and *σ* pairs than our model and can even take on negative values (Fig. 4).

**Figure 3.**
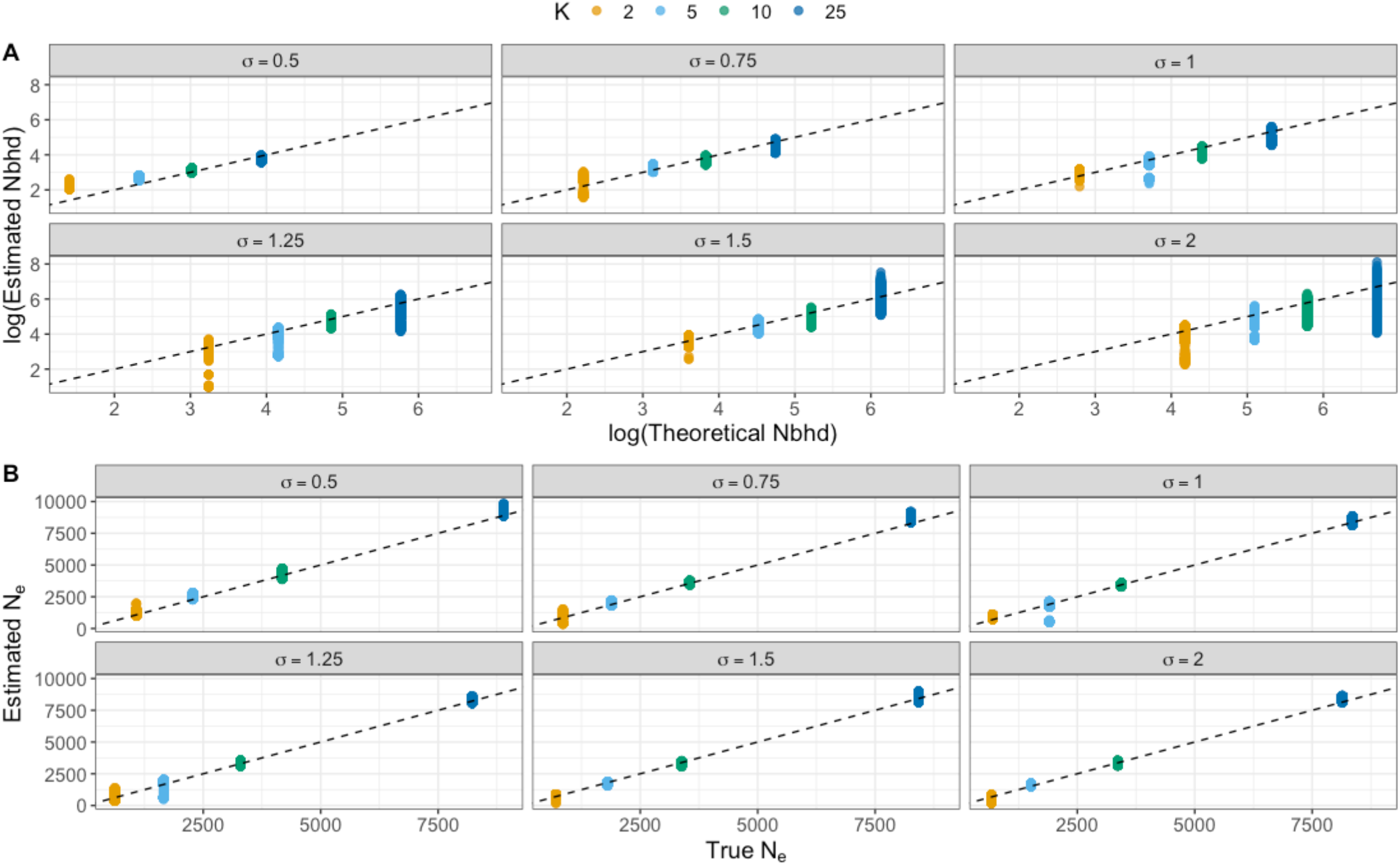
Model accuracy and precision. A) Estimated Wright’s neighborhood size (“Nbhd”) against the theoretical expectations along the 1:1 dashed line. B) Inferred *N_e_* from π_c_ against true *N_e_* estimated from the simulations (see text for details). Each panel represents simulated values of dispersal rate (*σ*), colors are different values of population density (*K*). Each point represents the posterior density from the best MCMC chain across all 10 simulation replicates.

**Figure 4.**
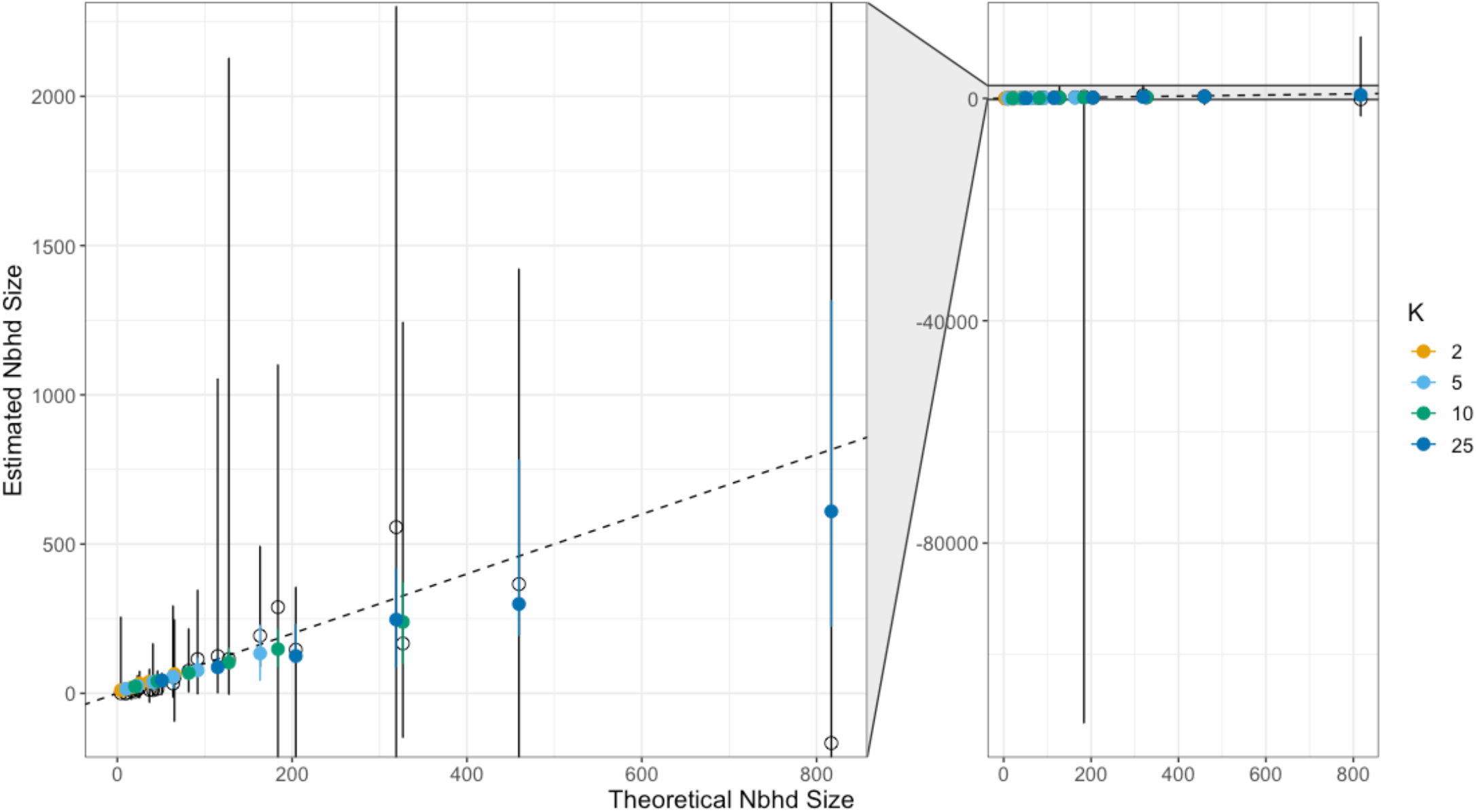
Comparison of the model presented in the text and Rousset’s method for estimating 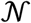. Black, open circles are the mean estimate using Rousset’s method and circles colored by *K* are the estimates under the model presented here. Vertical bars represent the 95% quantile, black for Rousset’s method and colored for the model in the text. The right panel shows the full range of Rousset’s 95% quantile for a pair of extreme values; the left panel is an inset capturing the narrower range of the 95% quantile of our estimates (colored bars within the longer black bars). The black dotted line represents the 1:1 relationship between the *x* and *y*.

The model also performed well at estimating π_c_. As noted above, the theoretical *N_e_* should scale negatively with increasing 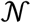 and *N_e_*, but positively with global census size and local density (regulated by *K*). While our model does not estimate *N_e_* directly, we estimate a proxy in π_c_; under neutrality, π_c_ ≈ 4*N_e_μ*. Rearranging, *N_e_* = π_c_ / 4*μ*, and we compare this value with the true *N_e_* estimated from *π*_*d*_*ij*_>20_ and the theoretical *N_e_* in Equation 9. When *N_e_* is estimated from *π*_*d*_*ij*_>20_, our model performs exceptionally well (Fig. 3B), generally falling along equality for all values of *σ* and *K*. When compared to the purely theoretical *N_e_*, our model underestimates *N_e_*, especially at higher values of *σ* and *K* (Fig. S2). We also compared estimated π_c_ across values of *σ* and *K* and found a general pattern of decreasing π_c_ at increasing *σ*, irrespective of population density (Fig. S1). This matches our expectations, based on theory (Wilkins 2004), that the total amount of genetic diversity in a finite, spatially structured population should increase with the degree of spatial structure, and therefore decrease with the scale of dispersal.

### Empirical results

For the Red Sea clownfish (*Amphiprion bicinctus*) dataset, our model estimated a 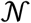 of 52.16 (95% CI: 50.54–54.04) (Fig. 5, S2). This neighborhood size is consistent with a recent localized population census study in the Gulf of Eilat, which found that the number of clownfish in the census year 2015 within a 200 × 50 m area was 52 fish (Howell et al. 2016). It is important to note that we have no independent information on the relevant spatial scale of dispersal or mating in Red Sea clownfish, and thus the correspondence between our estimates and the population census survey could be coincidental. We estimated a π_*c*_ of 0.003 and our estimated *m* (which is a compound parameter that includes long-distance dispersal and mutation) as 1.4e-9. Substituting *m* for the mutation rate, this produces an estimate of the long-term *N_e_* for clownfish as 535,714. It should be noted that while *A. bicinctus* is currently listed by the IUCN as “least concern” (Myers et al. 2017), several studies have suggested that clownfish are undergoing population declines, which could theoretically lead to a decoupling between 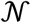 and *N_e_* (i.e., 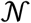 may reflect recent demographic shifts that *N_e_* has yet to be impacted by; Nanninga et al. 2015; Howell et al. 2016; Yosef et al. 2022).

**Figure 5.**
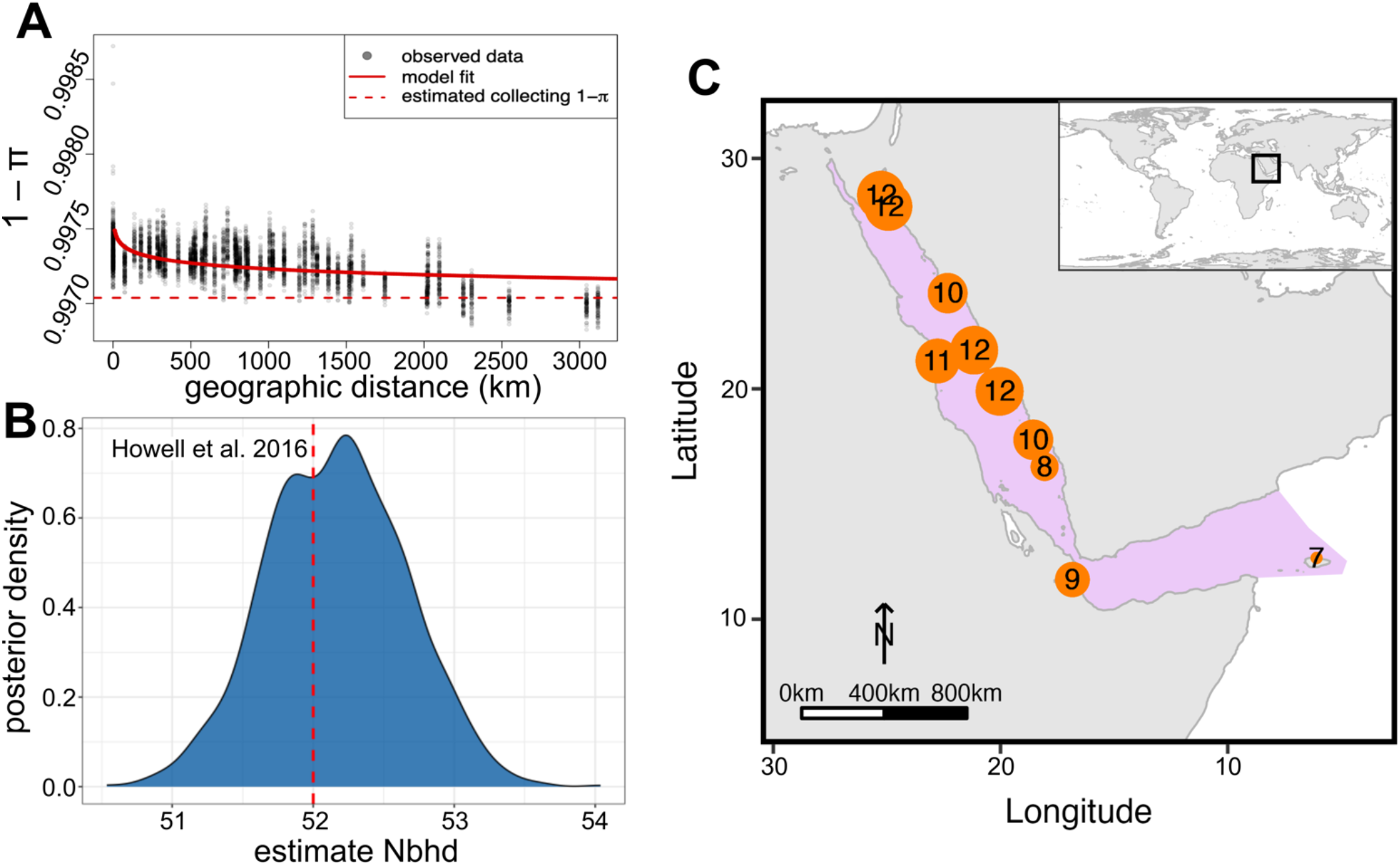
*Amphiprion bicinctus* dataset. A) Model fit, dark circles are the empirical observations and red line is the fit; dashed red line is estimated 1 – π_c_. B) Posterior density estimate for 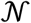 (“Nbhd”), with the red dashed line representing the number of individuals observed in the year 2015 in the Gulf of Eilat (Howell et al. 2016). C) Sample locations of *A. bicinctus*, with numbers representing sample size and circles scaled by sample size relative to other locations (Saenz-Agudelo et al. 2015).

## Discussion

An extensive historical literature exists on the biases that spatial structure introduces to commonly employed population genetic summary statistics (e.g., Bradburd & Ralph 2019; Battey et al. 2020). However, many studies still use measures of effective population size (such as Watterson’s *θ_w_* and *π*) that are naive to population structure (and the geographic sampling of individuals) in empirical systems that are spatially structured. Other attempts to develop statistical population genetic approaches that account for population structure by discretizing the habitat into demes that may be defined by sampling region or violation of Hardy-Weinberg (e.g. STRUCTURE – Pritchard 2000; Montana & Hoggart 2007), but in reality, many populations display a continuous pattern of isolation-by-distance that cannot be adequately reduced to a discrete stepping-stone model (Kimura & Weiss 1964). Indeed, even in a fine-scaled lattice population, Battey et al. (2020) showed that biases in estimates of Watterson’s θ and Tajima’s *D* emerge as the sample size per deme approaches the local effective size.

To investigate populations with continuous patterns of isolation-by-distance, we suggest that a single summary diversity statistic often does not capture the dynamics of interest and is skewed by the shape of the population genetic pedigree. For example, 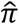 is dragged down by rapid coalescence in the scattering-phase but inflated by the influence of low dispersal in the collecting phase. Hence, 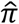 ceases to adequately reflect either process – it cannot tell us about local demography because it is upwardly inflated by deep-time coalescence, and it cannot tell us about ancient events because it is downwardly biased by local demography. Indeed, dispersal acts in opposing ways in these two phases: low dispersal causes local individuals to be more related on average and increases the length of the scattering phase, but over large distances and deeper timescales it inflates 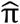 (Equation 9; Wilkins 2004). Our model generates estimates of a diversity statistic (π_c_) that is insensitive to the geography of sampling, and simultaneously provides estimates of Wright’s neighborhood size, thereby better capturing the spatial dynamics of the population.

Furthermore, our model is robust to a wide range of 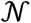 though the variance of our estimates of 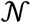 increases as the true 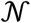 grows large. Theory and previous simulation-based research predicts that populations with very large 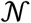 behave in a way that is approximately panmictic (Wright 1943), and populations need a considerably smaller 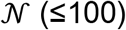 to differ substantially from random mating expectations (Battey et al. 2020). However, when subsampling the population randomly with respect to space, our simulated IBD curves indicate that 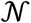 as low as ~200 generate results that superficially resemble panmixia. This finding is in line with theory from Wilkins (2002), who found that most of the population genetic pedigree occurs in the collecting phase. For IBD to be detectable at 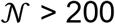, researchers would need to exhaustively sample local areas to ensure the collection of recent pedigree relatives due to how shallow the scattering phase is relative to the collecting.

One important caveat in interpreting the performance of our model on simulations is that there is not always a perfect correspondence between the quantities we are trying to estimate in our model (e.g., 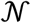 or π_c_) and parameter values we set in our simulations. For example, the Wright-Malécot IBD model assumes an infinite landscape, whereas our simulation model, in an effort to incorporate greater biological realism, has absorbing boundaries. We contend this is a more realistic depiction of species’ ranges, but it causes our simulation parameters to diverge from theory. Edge effects decrease both 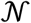 and *N_e_* relative to theoretical expectations because they reduce population density along the periphery of the range. Combined with lack of gene flow from even farther-flung regions, this reduction in diversity leads to neighboring individuals near edges being more related to one another on average than those sampled from the range center (Wilkins 2004; Rogan et al. 2023). Therefore, our model may be capturing this depression of 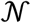 and *N_e_* relative to theoretical predictions, which may be especially apparent at high values of *K* and *σ*.

The other major way in which our simulation parameters diverge from their theoretical counterparts is in the difference between forward- and backward-time dynamics. The dispersal parameter we set in our SLiM simulations (*σ*) determines the midparent-offspring dispersal kernel in two dimensions forward-in-time, while the dispersal parameter in Equation 1, which affects both our estimates of 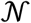 and π_c_, is the *effective* dispersal rate, and describes the dispersal kernel connecting offspring to their parents backwards-in-time (Cayuela et al 2018). The model of spatial competition and density-dependence (the implications of which are discussed more below) that we implement in our simulations causes the forward- and backward-time dispersal dynamics to differ. This discrepancy arises because, in our simulations, individuals are most likely to be born in regions of higher relative population density (due to the proximity required for parents to mate), where, if they remain, they will experience, on average, lower fitness. Therefore, individuals who disperse *farther* from their birth location are more likely to have higher fitness be represented in the population pedigree, which increases the effective dispersal rate relative to the forward-time rate we specify in our SLiM model. The impact of this discrepancy on the “true” value of 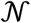 appears to be offset by the fact that effective population density is smaller than the value of K (for similar reasons) that we specify in our SLiM model. Nonetheless, it is helpful to be explicit about the differences between the meanings of the parameters we specify in our simulation model and that we estimate in our inference model.

### Model assumptions and shortcomings

While our model performs well on simulated data and the empirical dataset presented here, it makes several important assumptions that may often be violated in natural systems. Firstly, we assume that dispersal is random (within a specified dispersal kernel) and non-directional; thus, our model may poorly approximate patterns of isolation-by-distance in organisms that have directional movement, such as some marine planktonic species that may be carried by ocean currents or wind-dispersed pollen grains. Second, our model assumes that the habitat is relatively homogenous such that there are no major barriers to gene flow – i.e., patterns of isolation-by-distance are generated solely by the traversable Euclidean distance between two individuals. Strong physical or environmentally mediated barriers to dispersal may affect model performance (Ringbauer et al 2018, Wang & Bradburd 2014, Bradburd et al 2013). In a similar vein, our model ignores confounding factors such as local adaptation that may drive clinal patterns of relatedness (e.g., Pruisscher et al. 2018; Jofre & Rosenthal 2021).

As discussed in the Methods, the Wright-Malécot model relies on assumptions that are mutually incompatible (independent dispersal and homogeneous population density). In SLiM, we modeled populations with density-dependent selection to overcome Felsenstein’s “pain in the torus” (Felsenstein 1975) and maintain a roughly homogenous distribution of individuals across space. In this way, comparing the theoretical expectations from Wright-Malécot with a population model in which dispersal and density are not strictly independent is inexact. However, despite this discrepancy between theory and the simulated population, our model reasonably captured theoretical expectations, indicating that the Wright-Malécot formulation which underpins our model is robust to this violation.

Finally, our model relies on a user-defined *κ*, interpreted as the minimum-distance between two individuals in which the Wright-Malécot formulation breaks down and relatedness converges on 1 / 2*ρ* (Barton et al. 2002; Ringbauer et al. 2018). Preliminary exploration of the model indicates that model performance is poor at arbitrarily high *κ*, but not at low *κ*. This is due to the fact that, at higher *κ*, less of the scattering-phase is being captured by the model, leading to poor mixing. Ideally, *κ* would be estimated like the other parameters of the model; however, doing so has led to dramatic increases in computation time, and thus for the current work we opted to set a constant *κ*.

### Conclusions

For many organisms, geographical distance influences mate-choice, leading to patterns of continuous spatial structure. An important parameter governing the strength of isolation-by-distance is Wright’s neighborhood size 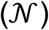, a theoretical quantity that describes the number of potential breeding individuals within a given dispersal radius. Previously, empirical researchers interested in estimating 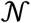 relied upon Rousset’s (1997; 2000) method, but this method requires either subsetting individuals into pseudopopulations (arbitrarily discretizing a potentially continuous reality) and calculating *F_ST_* between them or estimating pairwise *F_ST_* between individuals. The latter approach introduces a significant amount of noise into patterns of isolation-by-distance, and, in our simulation study, demonstrates poor performance due to high variance in results. Unlike Rousset’s estimator, our method shows good performance across values of 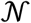, and offers the additional benefit of generating an estimate of the long-term effective population size (*N_e_*), which is linked to 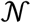 by dispersal. Here, we have presented a model that jointly estimates 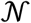 and long-term *N_e_* using an individual-based approach (i.e., it does not require arbitrary discretization via the lumping of samples). The introduced model produced reasonable estimates of the theoretical expectations from simulated data and performed well on an empirical dataset of clownfish. Future work will aim to generate an R package for ease of use for researchers (though code for performing the presented model is available in the Github link above). In addition, the model could be expanded to incorporate the potential for directional migration or patterns of isolation-by-environment (Wang & Bradburd 2014).

## Acknowledgments

We would like to thank members of the Bradburd lab – Leonard Jones, Meaghan Clark, Alex Lewanski, and Mike Grundler – as well as Teresa Pegan, John Wares, and Cynthia Riginos for helpful feedback on this manuscript. We are also grateful to feedback on developing this method from Luis Zaman and his lab, as well as Peter Ralph. This work was supported in part through computational resources and services provided by the Institute for Cyber-Enabled Research at Michigan State University. Research reported in this publication was supported by the National Institute of General Medical Sciences of the National Institutes of Health under Award Number R35GM137919 (awarded to G.S.B.). The content is solely the responsibility of the authors and does not necessarily represent the official views of the NIH.

## Data accessibility statement

Scripts for running the Wright-Malécot model and SLiM recipes are available at https://github.com/zachbhancock/WM_model.

## SUPPLEMENTARY MATERIALS

**Table S1.** Prior distributions for the model.

| Parameter | Distribution |
| --- | --- |
| $s$ | normal(0, 1) |
| $\log(\mu)$ | normal(-5, 1) |
| $\mathcal{N}$ | normal(100, 1000) |
| $\log(\gamma)$ | normal(0, 1) |
| $\log(\text{nugget})$ | normal(0, 1) |
| $\text{lik}(\text{pHom})$ | Wishart($L$, pHom) |

**Figure S1.**
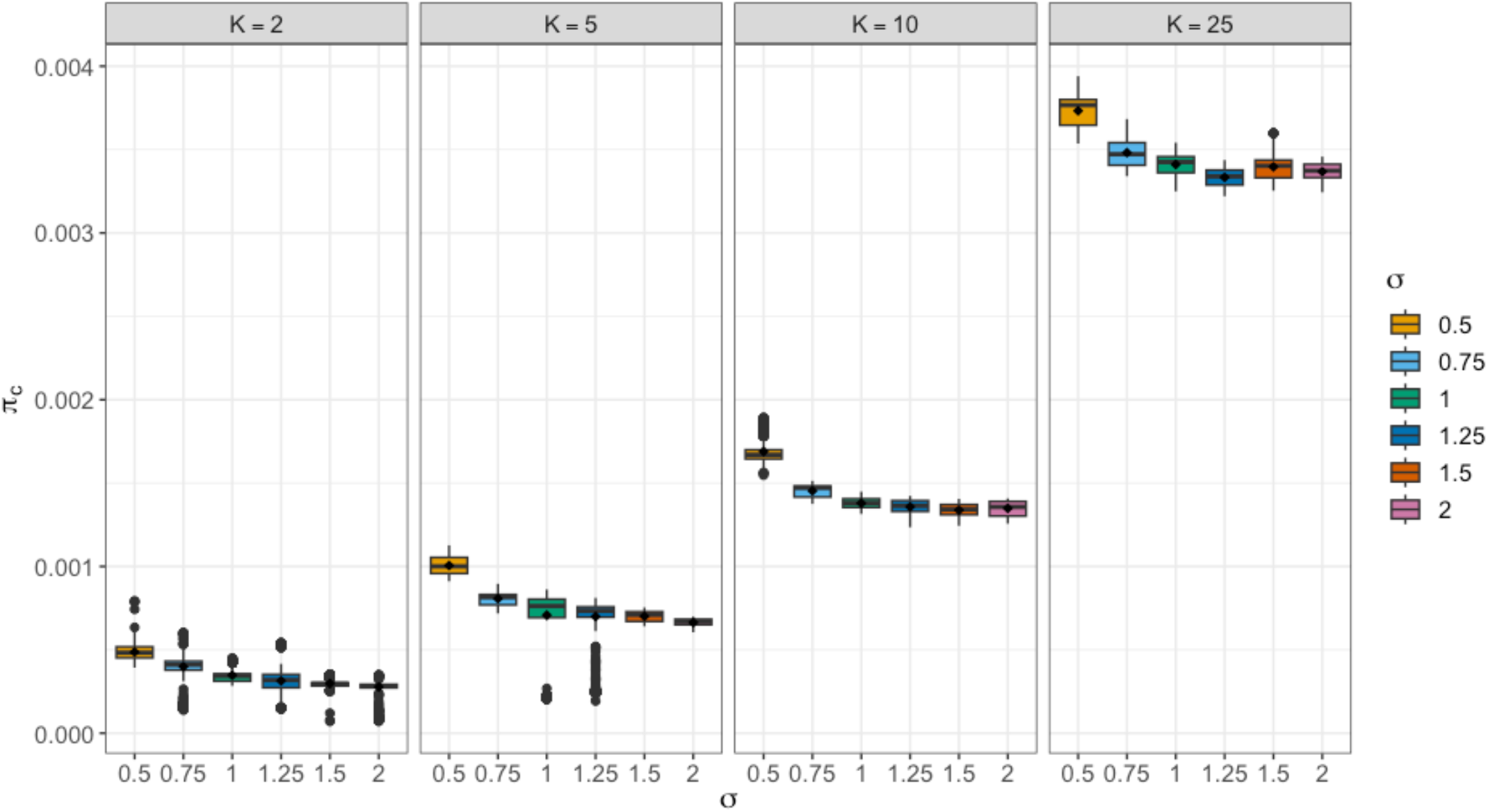
Estimated diversity during the collecting-phase (π_*c*_).

**Figure S2.**
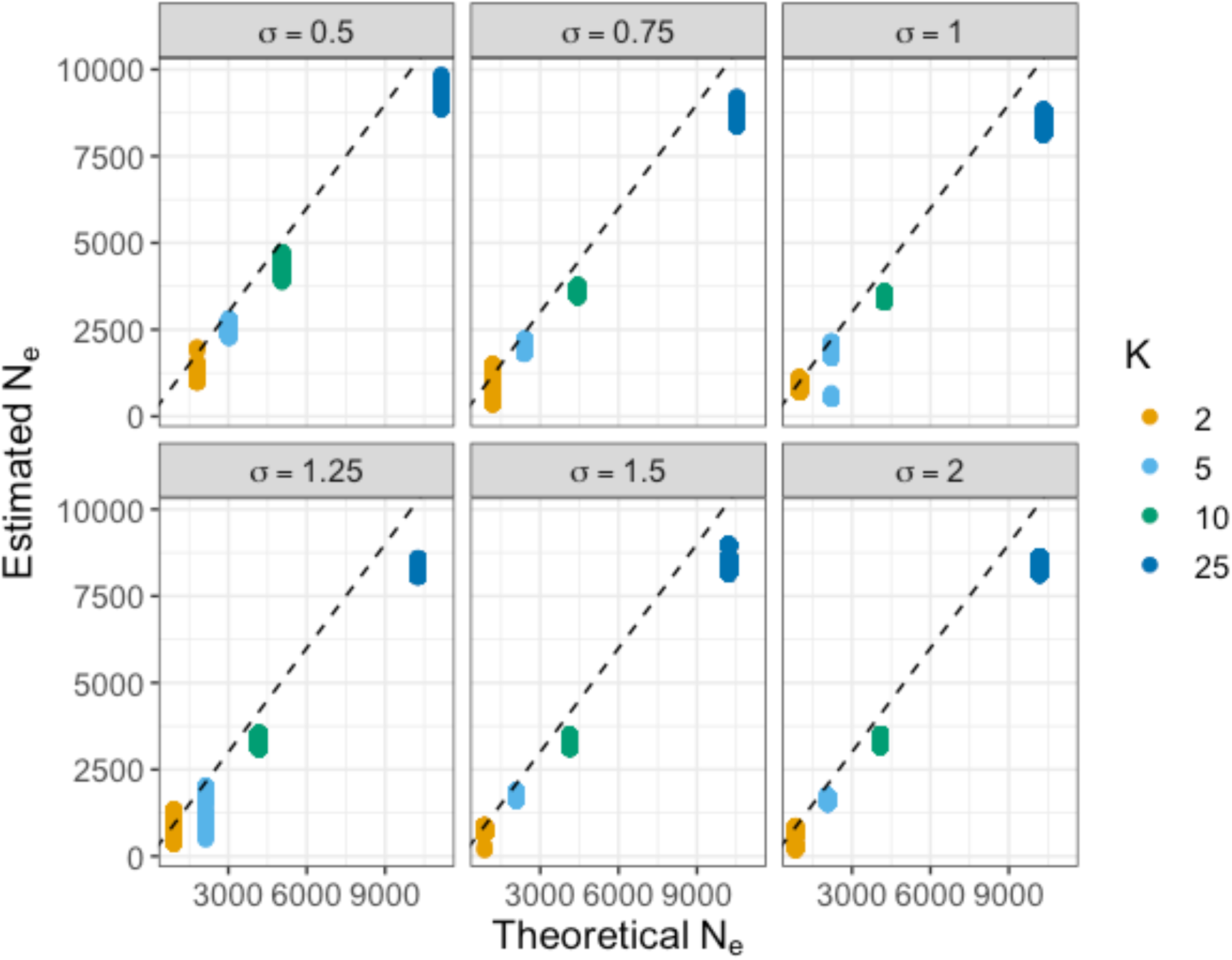
Estimated *N_e_* against theoretical *N_e_* from Eqn. 9. Unlike the results from Fig. 3B, we see the estimated *N_e_* is on average less than the theoretical; this is likely due to a mismatch between the simulated landscape that includes finite edges that varies population density across the range and the theoretical expectation of a homogenous distribution (see main text for further discussion).

**Figure S2.**
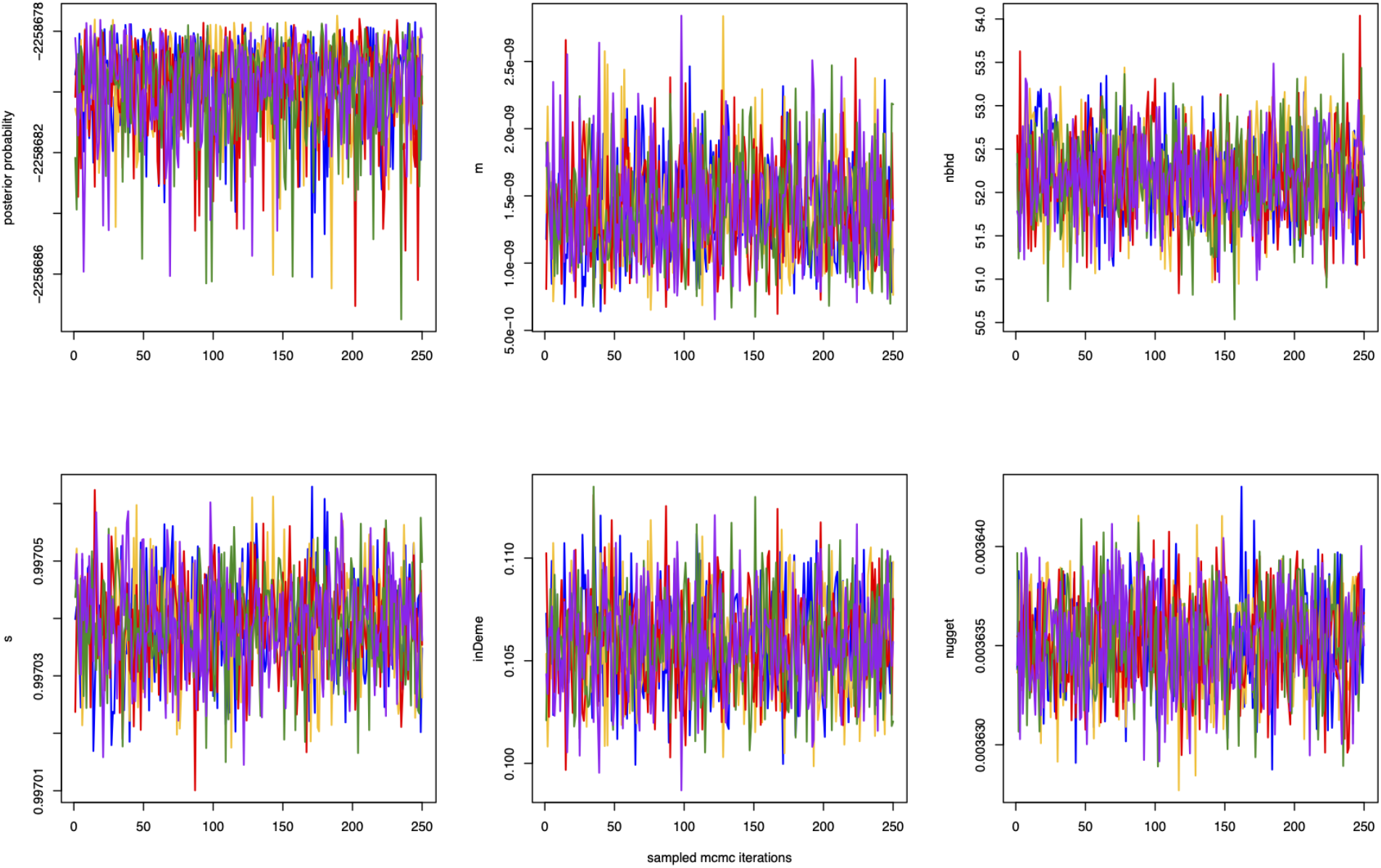
Trace plots of estimated parameter values for the *Amphiprion bicinctus* dataset; *m* is the compound mutation and long-distance migration rate; nbhd is Wright’s neighborhood size; *s* is the minimum relatedness between samples (see main text); “inDeme” is the coalescent rate for individuals within *κ* of one another; and the nugget is individual-level inbreeding.

**Figure S3.**
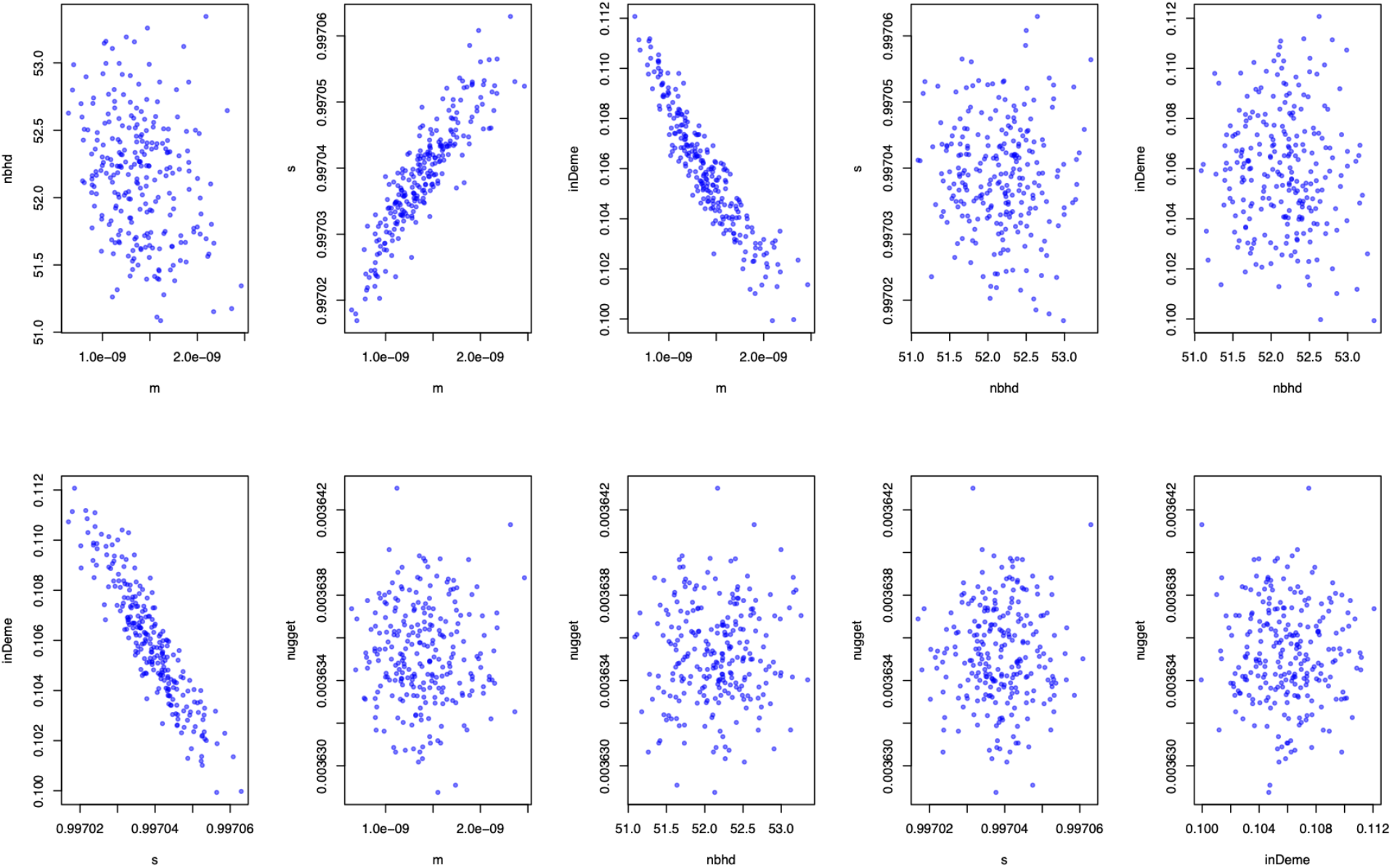
Autocorrelation plots between estimated parameter values for the *Amphiprion bicinctus* dataset; *m* is the compound mutation and long-distance migration rate; nbhd is Wright’s neighborhood size; *s* is the minimum relatedness between samples (see main text); “inDeme” is the coalescent rate for individuals within *κ* of one another; and the nugget is individual-level inbreeding.

